# PhoX – an IMAC-enrichable Crosslinking Reagent

**DOI:** 10.1101/556688

**Authors:** Barbara A. Steigenberger, Roland J. Pieters, Albert J.R. Heck, Richard A. Scheltema

**Affiliations:** Biomolecular Mass Spectrometry and Proteomics, Bijvoet Center for Biomolecular Research and Utrecht Institute for Pharmaceutical Sciences, Utrecht University, Padualaan 8, 3584 CH Utrecht, The Netherlands; Netherlands Proteomics Centre, Padualaan 8, 3584 CH Utrecht, The Netherlands; Department of Chemical Biology & Drug Discovery, Utrecht Institute for Pharmaceutical Sciences, Utrecht University, P.O. Box 80082, 3508 TB, Utrecht, The Netherlands

**Keywords:** XL-MS, IMAC, Tri-functional crosslinking reagent, Structural biology

## Abstract

Chemical crosslinking mass spectrometry is rapidly emerging as a prominent technique to study protein structures. Structural information is obtained by covalently connecting peptides in close proximity by small reagents and identifying the resulting peptide pairs by mass spectrometry. However, sub-stoichiometric reaction efficiencies render routine detection of crosslinked peptides problematic. Here we present a new tri-functional crosslinking reagent, termed PhoX, which is decorated with a stable phosphonic acid handle. This makes the crosslinked peptides amenable to the well-established IMAC enrichment. The handle allows for 300x enrichment efficiency and 97% specificity, dramatically reducing measurement time and improving data quality. We exemplify the approach on various model proteins and protein complexes, *e.g.* resulting in a structural model of the LRP1/RAP complex. PhoX is also applicable to whole cell lysates. When focusing the database search on ribosomal proteins, our first attempt resulted in 355 crosslinks, out-performing current efforts in less measurement time.

---

Crosslinking mass spectrometry (XL-MS) is a powerful tool that uses chemical reagents to investigate the structure of proteins and the complexes they form.^[1–5]^ The employed reagents are typically small bi-functional chemicals that covalently connect amino acids in close proximity. Most commonly highly efficient NHS-chemistry is used to capture the side chains of lysines in a protein.^[6–12]^ A spacer separates the reactive groups and as such, the crosslinking reagent acts as a distance constraint between the captured amino acids.^[13]^ After crosslinking the proteins in their native state, the protein sample is typically alkylated, reduced and finally digested into peptides by a protease, often trypsin. The full mixture of linear and crosslinked peptides is subsequently subjected to liquid chromatography tandem mass spectrometry (LC-MS/MS) for identification. After detection, crosslinked peptides provide valuable distance information on the protein tertiary structure in the form of intra-links (two peptides originating from the same protein) or on the protein quaternary structure in the form of inter-links (two peptides originating from different proteins).^[14]^ From these measurements, we and others found that the sought-for crosslinked peptides are completely overwhelmed by the linear peptides both in terms of numbers and abundance. To illustrate, from available data we estimate that the crosslink reaction efficiency is only 1-5% and, typically, relatively few lysine pairs are in close enough proximity to be crosslinked. Attempts to alleviate this situation have been made by extensive pre-fractionation of the peptide products using several chromatographic techniques.^[10,15–18]^ These techniques use properties that are exuberated for crosslinked peptides (*e.g.* size or charge), but are not unique. This typically results in large amounts of samples of which each still contains a high background of linear peptides. Other attempts integrate an enrichment handle directly on the crosslinking reagent resulting in a tri-functional reagent to separate crosslinked peptides from linear peptides. Two distinct approaches are typically used; the first one uses *e.g.* biotin as an enrichment handle on the spacer region.^[19–22]^ The second approach utilizes smaller functionalities on the spacer region like azides that allow performing bioorthogonal transformations after the crosslinking reaction. The crosslinked peptides are consequently functionalized by *e.g.* a biotin-handle using 1,3 dipolar cycloadditions (click-chemistry).^[23–25]^

The new strategy demonstrated here places a very small enrichable tag on the crosslinking reagent that possesses excellent enrichment properties in combination with the most efficient crosslinking chemistry. Inspired by the groundbreaking developments in phosphoproteomics, where immobilized metal affinity chromatography (IMAC) has reached levels of > 95% enrichment specificity,^[26]^ we hypothesized that a phosphate-group could be an ideal enrichment handle. Such a handle has the additional advantage that IMAC enrichment is nowadays a nearly routine protocol in most proteomics laboratories,^[27–29]^ and high levels of automation with liquid handling platforms can be achieved.^[30]^ The lability of the phosphate-group during sample handling and mass spectrometric acquisition makes detection however problematic.^[31,32]^

We tackled this issue by altering the phosphate-group to a phosphonic acid, replacing the labile P-O bond by a stable P-C bond. Such a phosphonic acid handle is still amenable to IMAC enrichment and thus PhoX enables a completely novel and highly efficient enrichment approach for the field of XL-MS. We demonstrate the versatility of PhoX on various proteins and protein-complexes and even a full cell lysate (see Supplementary Notes 1-4 for further details on sample preparation, enrichment, data acquisition and data analysis).

## Synthesis

In the design of PhoX we attempted to minimize the spacer-length, and optimize the hydrophobicity of the reagent to ensure maximum reaction efficiency (see Supplementary Note 5). We make use of an aromatic core structure as the vehicle for two NHS esters and the enrichable phosphonic acid (compound **5**; see scheme in Fig. 1a). This results in a planar rigid spacer providing a fixed length of 5 Å, leading to a maximum distance constraint of about 20 Å when including the flexible lysines side-chains. A major hurdle in the synthesis of such a reagent lies in the combination of NHS-ester and phosphonic acid functionalities. Carboxylic acids, which are the common precursors for NHS-esters, as well as phosphonic acids, can be activated by carbodiimide reagents, leading to a mixture of undesired NHS-ester functionalities attached to the phosphonic acid moiety. To counter this, the phosphonic acid was protected by a dibenzyl-ester during the NHS-ester synthesis. Dibenzyl-esters can be selectively removed by hydrogenation with palladium on carbon (Pd/C), leaving the NHS esters intact.

**Figure 1.**
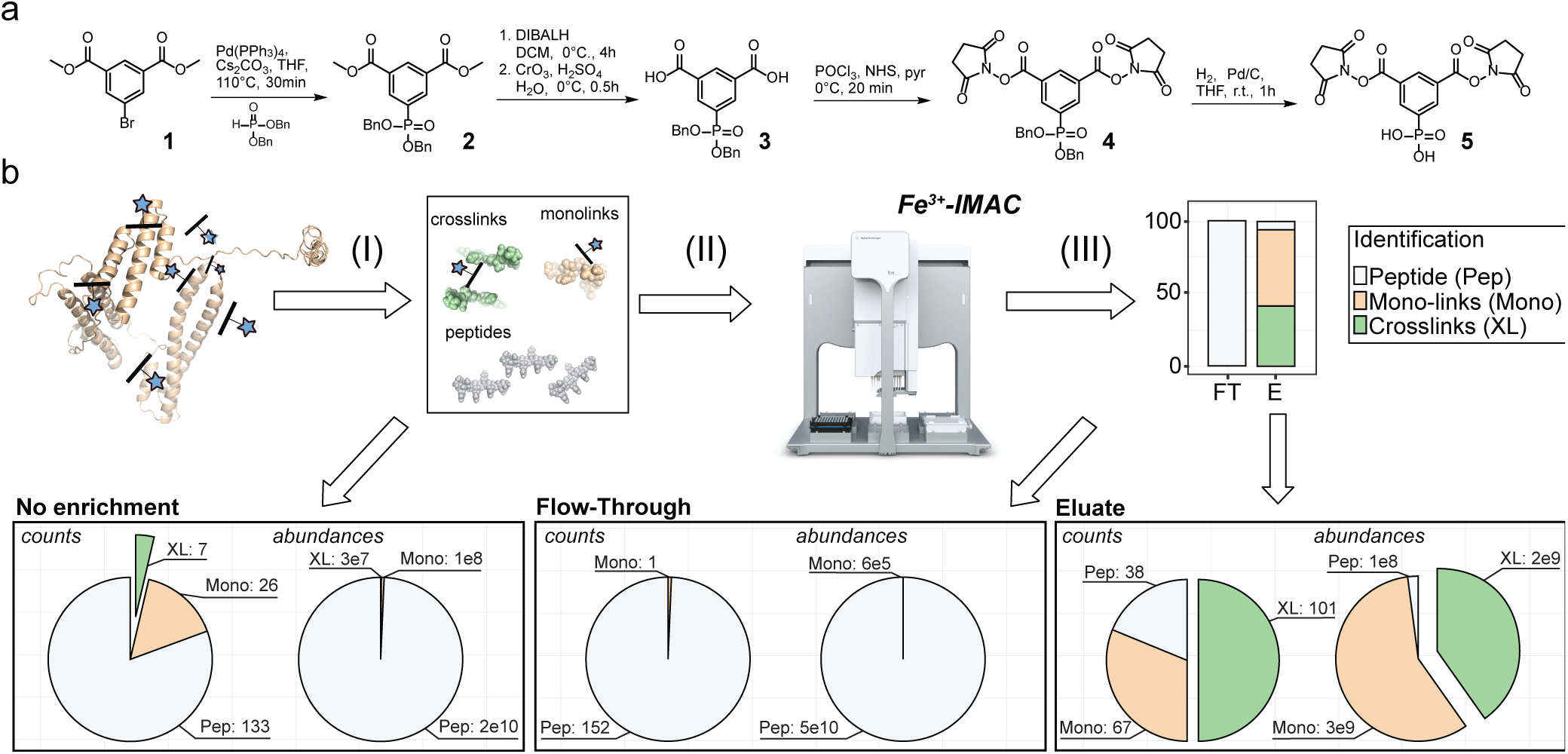
PhoX synthesis, workflow and performance. **(a)** The synthesis of PhoX can be achieved in five steps starting from commercially available precursors. **(b)** After crosslinking the proteins in their native state, the proteins are denatured, reduced, alkylated and digested to peptides (I). The mixture of peptides contains 3 different kinds of products: unmodified peptides (grey), monolinks (orange) and crosslinks (green) (II). Crosslinked and monolinked peptides are enriched using Fe-IMAC on a liquid sample handling platform providing high sample throughput (III). Direct measurement of the crosslinked peptides produces low amounts of crosslink identifications (left panel), while measurement following Fe-IMAC enrichment result in many crosslink identifications in the eluate (middle panel) and none in the flow-through (right panel).

The final crosslinking reagent **5** was synthesized using the commercially available dimethyl 5-bromoisophthalate **1** as starting compound, which acts as our spacer (see Fig. 1a). Compound **1** was subjected to a palladium-catalyzed cross-coupling reaction to form a C-P-bond in the coupling product **2**. As deprotection of the methyl ester of **2** under basic conditions yielded partly the free phosphonic acid, we reduced the methyl ester by treatment with diisobutylaluminium hydride at 0 °C to the alcohols. The di-alcohol was oxidized in quantitative yields to the corresponding di-carboxylic acid **3** with the Jones reagent. No by-products were observed using this reaction and only a crude work-up was necessary. Activation of compound **3** with phosphoroxychlorid in pyridine at 0 °C and subsequent addition of N-hydroxysuccinimide yielded the NHS-ester functionalized reagent **4** within 20 min. As the final step, the dibenzyl phosphonic acid **4** was subjected to a hydrogenation reaction using Pd/C to yield the free phosphonic acid while keeping the NHS esters intact. After filtering of the Pd/C the crosslinking reagent **5** could be used without further purification.

## Performance on model proteins

Verification by tandem mass spectrometry showed that, in contrast to the P-O bond, the P-C bond is indeed very stable (see Supplementary Note 6). To visualize the performance of our reagent, we crosslinked human hemoglobin and ran the crosslinked protein on SDS-page. We tested a range between 0.5 and 5 mM for our crosslinking reagent and benchmarked it against the commonly used crosslinking reagent disuccinimidyl suberate (DSS). Based on the SDS-PAGE we estimated for both linkers that the optimal concentration is 1 to 2 mM, albeit that DSS appears a little more reactive which can easily be compensated with a higher crosslinker concentration. For further performance tests we used BSA, a protein which has emerged as a standard for testing crosslinking performance.^[33–35]^ Crosslinking with 1 mM of PhoX and in-solution digesting the sample after crosslinking provided the input for our enrichment (see Fig. 1b; Supplementary Note 7). We retained part of the input as control (*i.e.* non-enriched) prior to enrichment with Fe-IMAC and collected both the flow-through as well as the eluate. The control resulted in the identification of 7 crosslinked peptides, 26 mono-links (one NHS-ester of the reagent reacted with the peptide and the other is hydrolyzed, providing no structural information) and 133 linear peptides. The abundance levels of the crosslinked peptides – constituting 0.14% of total ion intensity – are not even visible in the high background levels of linear peptides and as such will be difficult to analyze (Fig. 1b; left panel). We retrieved from the flow-through no crosslinked peptides, 1 mono-link and 152 linear peptides. As expected, investigation of the abundances shows the same general trend as the control (Fig. 1b; middle panel). The fact that the flow-through contained no crosslinked peptides indicates that the crosslinked peptides were successfully bound to the IMAC material. In sharp contrast, we retrieved from the eluate 101 crosslinked peptides, 67 mono-linked peptides and 38 linear peptides. Investigation of the abundance values showed a radically different picture from the control (Fig. 1b; right panel). Even though there are more crosslinked peptides than mono-links, mono-links show a 1.3 x higher intensity as it is easier to form a mono-link than a crosslink and both bind to the IMAC material. Calculation of enrichment efficiency and specificity from the detected abundance levels demonstrates a 300x enrichment of crosslinked peptides, and a 97% enrichment specificity. Mapping the detected crosslinked peptides on the structure of BSA we found 60% were within the 20 Å distance limit we set for this linker (Supplementary Note 8). Given that our sample is crosslinked in-solution some crosslinks may result from the lower abundant dimer and trimer oligomers of the BSA protein, as also observed previously.^[36]^

To assess the performance of PhoX crosslinking in a protein environment of similar complexity as those typically encountered for purified protein complexes, we applied the crosslinker on the Pierce Intact Protein Standard Mix consisting of six proteins (Supplementary Note 9). As these proteins do not naturally interact no inter-links are expected. After optimization of the PhoX crosslink reagent concentration to 2 mM we detected after IMAC enrichment 134 crosslinks. We found two inter-links between DNA Polymerase I and Thioredoxin as well as DNA Polymerase I and Protein G, which are most likely false positives. An additional 5 inter-links are reported in the table, which all derive from Protein A and G and should actually be considered intra-links as the sample contains the fusion protein Protein A/G. A similar analysis of the flow-through revealed no crosslinked peptides, demonstrating the effectiveness of the enrichment approach also in analyzing this more complex protein mixture.

## Enrichment efficiency

To further test the efficiency of our approach for enriching low abundant crosslinked peptides, we next spiked PhoX-crosslinked BSA peptides into a background of an E. coli tryptic digest in a broad dilution range. We chose the E. coli digest as background, as it contains relatively low levels of potentially co-enriched phosphorylated peptides.^[37]^ Such a dilution range can be done in two ways, each representing a different outcome (Table 1; Supplementary Note 10). The first approach consists of mixing a decreasing amount of PhoX-crosslinked BSA peptides in a fixed background of E. coli tryptic peptides – we term this approach ‘Sensitivity Test’. This mimics the concentration levels at which crosslinks can still be recovered from a full lysate using our approach. For this, a concentration of crosslinked BSA peptides ranging from 16 *µ*g down to 0.5 *µ*g was spiked into a background of 100 *µ*g of E. coli tryptic peptides. Given that we observed that only 0.14% of the total peptide weight was represented by crosslinked peptides, we estimate at its highest dilution point crosslinked peptides represent approximately 700 pg. These detectable pico-gram levels are within the range typically achieved by multiple reaction monitoring (MRM)^[38]^, although our measurements do not make use of a targeted approach. MS analysis of the enriched samples reveals that crosslinked peptides are reliably identified at approximately 10 ng levels with 56 identified crosslinks (93% of undiluted crosslinked BSA – slightly higher compared to the amount detected for the recovery experiment which we attribute to experimental variation); although at the lowest level of 700 pg still two crosslinks can be identified.

**Table 1.** Setup and results for the spike-in experiments.

| Sensitivity |  |  | Recovery |  |  |
| --- | --- | --- | --- | --- | --- |
| Ratio BSA:*e. coli* | Final XLs (ng) | XL count | Ratio BSA:*e. coli* | Final XLs (ng) | XL count |
| 16:100 | 22.4 | 69 | 10:0 | 14 | 36 |
| 8:100 | 11.2 | 60 | 10:10 | 14 | 30 |
| 4:100 | 5.6 | 40 | 10:100 | 14 | 22 |
| 2:100 | 2.8 | 29 | 10:1000 | 14 | 21 |
| 1:100 | 1.4 | 15 | - | - | - |
| 0.5:100 | 0.7 | 12 | - | - | - |

The second approach consists of mixing equal amounts of crosslinked BSA peptides in an increasing background of E. coli peptides – we term this approach ‘Recovery Test’ – for which we used 10 *µ*g of crosslinked BSA. This ensures that an equal amount of crosslinked peptides can be recovered and measured; removing effects of low concentration prohibiting successful detection by LC-MS/MS. Here we estimate that the crosslinked peptides represent approximately 14 ng. Approximately 30 crosslinked peptides at these concentration levels are recovered efficiently without much loss, indicating that the enrichment is not significantly affected by high background levels of linear peptides.

## Elucidation of LRP1-RAP binding

Its enrichable nature, small footprint and short spacer makes PhoX attractive for the elucidation of interfaces between tightly interacting proteins. We demonstrate this here investigating the interaction of the lipoprotein receptor-related protein 1 (LRP1; Fig. 2a) and the receptor associated protein (RAP; Fig. 2b); further information on these proteins is provided in Supplementary Note 11. It was biochemically shown that RAP, which comprises of three domains (D1-3), has a high affinity for Cluster II.[39] D3 tightly interacts via exposed lysines (K293 and K307) to acidic pockets exposed on the surface of LRP1 (D283 and D326). The D2 domain exhibits lower affinity binding, while the separate D1 domain showed no binding. Nevertheless, it is so far unknown how full-length RAP binds to LRP1, despite previous attempts.^[40]^ Here we investigate the binding of full-length RAP to Cluster II of LRP1 with PhoX (see Supplementary Note 11 for information on the structural modelling). Using PhoX we find 31 intra-links on RAP, 3 intra-links on LRP1 and 16 inter-links between the two proteins. Investigation of the intra-links within the individual domains of RAP revealed that the observed distance constraints satisfy the maximum length of PhoX, indicating that the experiment successfully yielded useful structural information. Investigation of the inter-domain crosslinks revealed a considerably different picture where almost all distances are outside the range (on average 37.3±20.6 Å) indicating that RAP under-goes extensive structural rearrangement upon binding LRP1 (Fig. 2c). Notably, based on NMR data the possibility of such rearrangement had already been predicted.^[41]^

**Figure 2.**
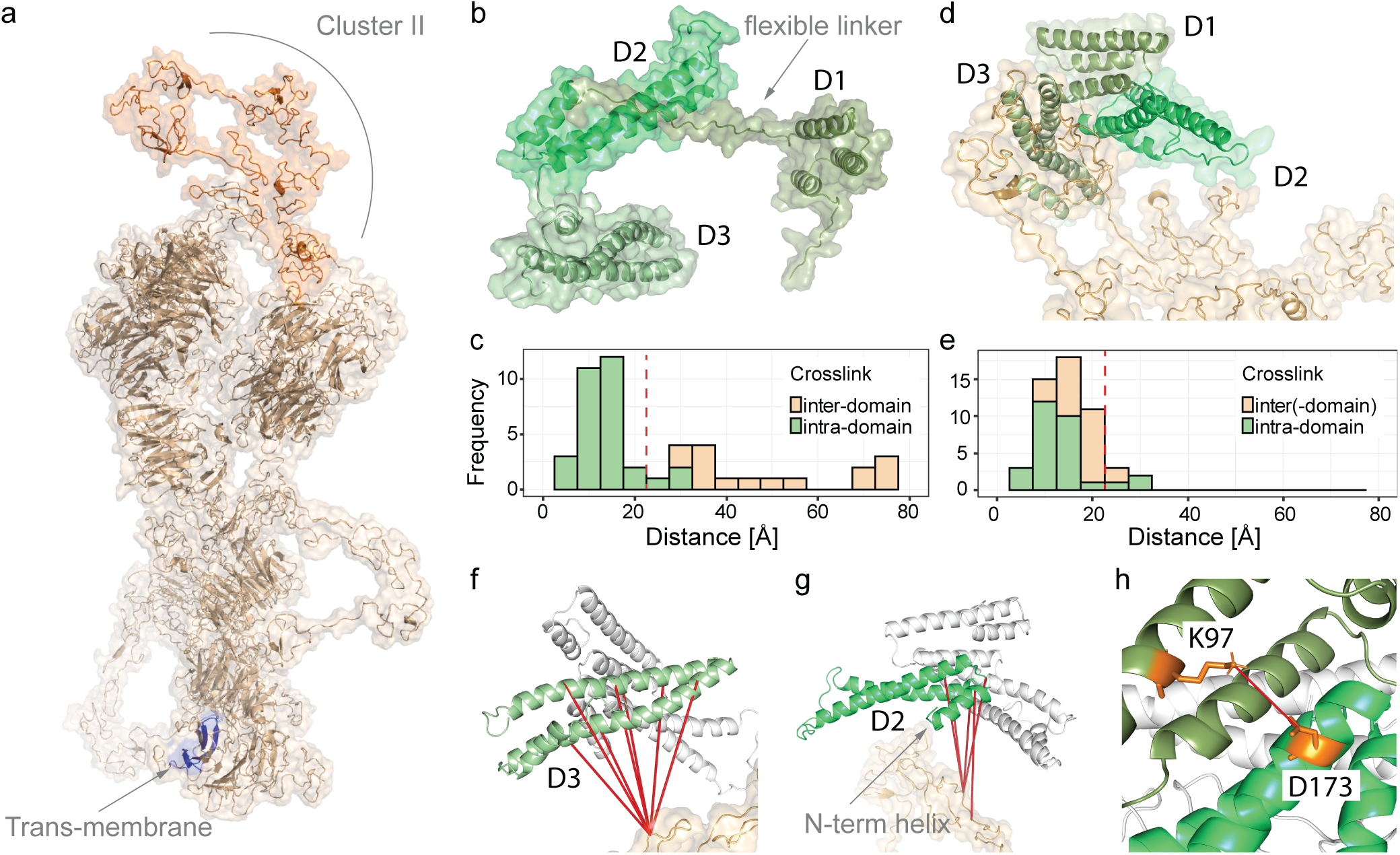
Application of PhoX to investigate the binding of full-length RAP to LRP1 Cluster II. **(a)** Predicted structural model for full length LRP1, with Cluster II (PDB: 1n7d) and the trans-membrane region highlighted. **(b)** Structure of RAP with the individual domains indicated (predicted from PDB: 2p03). **(c)** Measured C*α*-C*α* distances for intra- and inter-domain crosslinks on the structure of RAP (maximum distance constraint indicated with red, dashed line). **(d)** Multi-body docking of the individual domains of RAP to LRP1 using HADDOCK. **(e)** Measured C*α*-C*α* distances after docking (maximum distance constraint indicated with red, dashed line). **(f)** Crosslinks between the D3 domain of RAP and LRP1. **(g)** Crosslinks between the D2 domain of RAP and LRP1. **(h)** Domain D1 (K97) in close proximity to domain D2 (D173).

To further investigate this major structural rearrangement, we performed multi-body docking to LRP1 using HADDOCK.^[42]^ For this, the individual domains of RAP were separated from each other by removing the long linker regions. The resulting best scoring structural model, which retains the connectivity of the lysines and the acidic pockets intact, was selected for further analysis (Fig. 2d). This model places K293 of RAP in the acidic pocket around D326 of LRP1 and K307 of RAP in the acidic pocket around D283 of LRP1. In this model, the domains D2 and D3 connect with LRP1, while the domain D1 exclusively interacts with the D2 and D3 domains. Inspection of the measured distances shows that over 90% of the crosslinks in this model are within the set distance constraints (Fig. 2e). Inspection of the crosslinks between LRP1 and the D3 domain of RAP shows that crosslinks are detected over the full length of this domain (Fig. 2f). This can readily be explained by the strong interactions of the two lysines on D3 for which no crosslinks were found, enforcing a close interaction of the full length of this domain to LRP1. In contrast, crosslinks between LRP1 and the D2 domain of RAP are located on the short N-terminal helix of this domain lifting a large portion of this domain of the surface of LRP1 (Fig. 2g). This suggests that this helix is interacting solely with LRP1, previously suggested based on its high sequence conservation.^[39]^ The placement of the D1 domain suggests a role for this domain to lock the domain D2 into place upon LRP1 binding. Structurally a connection could potentially be made through a salt bridge between D173 and K97, a lysine which previously was shown to be shielded upon binding to LRP1^[44]^ and for which we observe no crosslinks.

## Application to full human lysate

Advanced mass spectrometry in combination with crosslinking has the potential to unbiasedly map protein-protein interaction networks *in situ* of the cell. Major hurdles associated to the low stoichiometry of the crosslinking reaction, the presence of two peptides during fragmentation, and the overwhelming complexity for database searches however still remain. Typically, in such an experiment extensive fractionation has to be performed to reduce the complexity to successfully capture crosslinked peptides – an area where PhoX could provide an excellent alternative. We tested the enrichment power of PhoX on a natively lysed human substrate without fractionation. As PhoX is not (yet) a MS-cleavable crosslinker, database searches against a whole human proteome are still a nearly unsurmountable problem. To tackle this, we chose to search only for ribosomal associated proteins, as they are abundantly present and known to be associated, allowing us to use a reduced FASTA file of only 438 human ribosomal proteins for the search in Proteome Discoverer with XlinkX integrated.^[7,14]^ From a single 180 min LC-MS/MS run on the IMAC eluate we could identify readily 307 crosslinks (from 684 spectra) on ribo-some associated proteins. As enrichment of crosslinks by Fe-IMAC also involves simultaneous enrichment of phosphorylated peptides, we envisioned that de-phosphorylation of peptides prior to enrichment could further improve this result. Whereas phosphate-esters present on phosphorylated peptides are hydrolyzed upon treatment with phosphatase treatment, the carbon-phosphor bond present on the phosphonic acid group of the crosslinking reagent will not be affected. After enrichment of CIP phosphatase treated crosslinked lysate, we could increase the amount of crosslinks detected on ribosomal associated proteins to 355 (from 787 spectra), an improvement of 15%. Further indication that the phosphatase treatment was successful came from the observed drop of 1120 to 59 PSMs for phosphorylated peptides in the non-treated lysate versus treated lysate. In this dataset, 158 of the 307 crosslinks (identified by 314 spectra) are solely on the 80S ribosome, a 50% higher number than reported in previous reports.^[7,45]^ Of the 158 crosslinks, 71 crosslinks could be mapped on the cryo-EM structure of the 80S human ribosome (PDB:4v6x), the others are predominantly in regions not resolved in the structural model of the ribosome. Of the mapped crosslinks, 90% are within the defined distance constraint of 20 Å (Supplementary Note 12). Of interest though, is that these numbers were obtained with a single shot measurement of 180 min. For previous efforts, extensive fractionation was required to get similar numbers (*e.g.* for the XL-MS study on the nucleus, 86 measurements, totaling 172 hrs of measurement time was needed to extract 87 crosslinks on the ribosome^[45]^), clearly demonstrating the sensitivity and efficiency of PhoX.

## Conclusion

Here we presented a novel enrichable crosslinking reagent functionalized with a phosphonic acid group on the spacer region termed PhoX (full details on the synthesis are provided in Supplementary Note 13). PhoX integrates a small phosphonic acid functionality on the spacer region which is amenable to large scale and efficient IMAC enrichment as commonly used in phosphoproteomics. The linker however incorporates a stable C-P bond ensuring permanent decoration of the crosslinked peptides with the enrichment handle. IMAC is not only a unique approach for the enrichment of crosslinked peptides, but also allows for a best-case 300x enrichment efficiency and 97% enrichment specificity according to our data. The described workflows in combination with PhoX, we expect, will simplify XL-MS approaches on several fronts:

- Efficient crosslinking with a small, non-sterically hindered crosslinking reagent;
- A condensed spacer length facilitating higher precision for modelling protein structures;
- A simplified and fast workflow, which uses standardized phospho-proteomics techniques for the enrichment of crosslinked peptides. These techniques have been implemented in many proteomics laboratories, facilitating adoption of the reagent. Competing molecules for the IMAC enrichment such as phospho-peptides and nucleic acids can easily be removed, while the much more stable phosphonic acid moiety for the enrichment of crosslinked peptides remains intact;
- Fe-IMAC for the enrichment of crosslinked peptides enables high-throughput sampling in a 96-well plate format;
- LC-MS measurement time can be decreased as one fraction is sufficient to achieve in-depth crosslink identification. In those cases where further fractionation is required, PhoX is highly compatible with orthogonal approaches like high-pH fractionation;
- Removal of linear peptides enables reliable identification of crosslinked peptides resulting in high quality fragmentation spectra for crosslinked peptides.

As always, there remains room for improvement. Likely, next versions of PhoX will come with different reactive groups, incorporate stable isotopes and gas-phase cleavable moieties. Still, we envision already widespread adoption of XL-MS supported by this first version of PhoX, as it already tremendously facilitates in-depth studies into the dynamics of proteins, protein complexes and interactomes; diminishing analysis time and sensitivity by sizeable factors.

## Acknowledgments

We thank all members of the Heck-group for their helpful contributions, Dr. Domenico Fasci for providing the cell lysate and Prof. Dr. Sander Meijer and Carmen van der Zwaan from Sanquin for the kind gift of the LRP1 Cluster II and RAP proteins. We acknowledge financial support by the large-scale proteomics facility Proteins@Work (Project 184.032.201) embedded in the Netherlands Proteomics Centre and supported by the Netherlands Organization for Scientific Research (NWO). Additional support came through the European Union Horizon 2020 program FET-OPEN project MSmed (Project 686547), the European Union Horizon 2020 program INFRAIA project Epic-XS (Project 823839), and a seed grant kindly provided by the Utrecht Institute for Pharmaceutical Sciences (UIPS).

## Contributions

R.A.S. and A.J.R.H. conceived of the study. R.J.P. supported the synthesis and provided input for the design of the reagent. B.S. developed and synthesized the reagents and performed the experiments. B.S. and R.A.S. performed data analysis. All authors critically read and edited the manuscript.

